# The mod-minimizer: a simple and efficient sampling algorithm for long *k*-mers

**DOI:** 10.1101/2024.05.25.595898

**Authors:** Ragnar Groot Koerkamp, Giulio Ermanno Pibiri

## Abstract

**Motivation:** Given a string *S*, a *minimizer* scheme is an algorithm defined by a triple (*k, w*, 𝒪) that samples a subset of *k*-mers (*k*-long substrings) from a string *S*. Specifically, it samples the minimal *k*-mer according to the order 𝒪 from each window of *w* consecutive *k*-mers in *S*. Because consecutive windows can sample the same *k*-mer, the set of the sampled *k*-mers is typically much smaller than *S*. More generally, we consider substring sampling algorithms that respect a *window guarantee*: at least one *k*-mer must be sampled from every window of *w* consecutive *k*-mers. As a sampled *k*-mer is uniquely identified by its absolute position in *S*, we can define the *density* of a sampling algorithm as the fraction of distinct sampled positions. Good methods have low density which, by respecting the window guarantee, is lower bounded by 1*/w*. It is however difficult to design a sequence-agnostic algorithm with provably optimal density. In practice, the order 𝒪 is usually implemented using a pseudo-random hash function to obtain the so-called *random* minimizer. This scheme is simple to implement, very fast to compute even in streaming fashion, and easy to analyze. However, its density is almost a factor of 2 away from the lower bound for large windows.

**Methods:** In this work we introduce *mod-sampling*, a two-step sampling algorithm to obtain new minimizer schemes. Given a (small) parameter *t*, the mod-sampling algorithm finds the position *i* of the minimal *t*-mer in a window. It then samples the *k*-mer at position *i* mod *w*. The *lr-minimizer* uses *t* = *k* − *w* and the *mod-minimizer* uses *t* ≡ *k* (mod *w*).

**Results:** These new schemes have provably lower density than random minimizers and other schemes when *k* is large compared to *w*, while being as fast to compute. Importantly, the mod-minimizer achieves optimal density when *k* → ∞. Although the mod-minimizer is not the first method to achieve optimal density for large *k*, its proof of optimality is simpler than previous work.

We provide pseudocode for a number of other methods and compare to them. In practice, the mod-minimizer has considerably lower density than the random minimizer and other state-of-the-art methods, like closed syncmers and miniception, when *k > w*. We plugged the mod-minimizer into SSHash, a *k*-mer dictionary based on minimizers. For default parameters (*w, k*) = (11, 21), space usage decreases by 15% when indexing the whole human genome (GRCh38), while maintaining its fast query time.

## 1 Introduction

The concept of *minimizer* was simultaneously introduced by Roberts et al. [23] and Schleimer et al. [26] as an algorithm that samples a small set of *k*-mers from a string *S*. More precisely, a minimizer scheme is defined by a triple (*k, w*, 𝒪) and works as follows. In each window of *w* consecutive *k*-mers of *S*, the minimal *k*-mer according to the order 𝒪 is chosen. The collection of distinct (according to their position in *S*) sampled *k*-mers represents a succinct sketch of *S*, as a minimizer scheme tends to choose the same *k*-mer across adjacent windows. This property makes the minimizer highly versatile, facilitating memory reduction and faster processing in various bioinformatics applications such as sequence comparison [25], assembly [5], construction of compacted De Bruijn graphs [2, 3], and sequence indexing [18, 19, 12, 7, 20].

More generally, the effectiveness of a *sampling algorithm* is measured by its ability to sample a small number of *k*-mers from *S*, while ensuring a *window guarantee*, that is, at least one *k*-mer is sampled from every window. Since a *k*-mer can be uniquely identified with its absolute position in *S*, we informally define the *density* of a sampling algorithm as the ratio between the number of distinct sampled positions and the length of *S*. The lower the density of a scheme, the better the scheme, as it improves space and time resources for the aforementioned applications.

The necessity to maintain the window guarantee implies a lower bound on density, which is 1*/w*. Achieving optimal performance, i.e., density close to 1*/w*, is challenging, especially when the scheme is required to be sequence-agnostic. In fact, practitioners usually resort to implementing the order 𝒪 using a pseudo-random hash function to obtain what is known as the *random* minimizer scheme. While this scheme is straightforward to implement, computationally efficient (even in streaming scenarios), and analytically easy to analyze, its density falls short of the theoretical lower bound by almost a factor of 2 for large *w*.

Previous research has thus predominantly focused on developing methods that achieve lower density compared to the random minimizer, both theoretically [11] and practically [4, 17, 27]. However, these methods either pose challenges in terms of analysis and intuitive understanding [11, 27], or are computationally expensive [17].

### Contributions

In this work we introduce *mod-sampling*, a novel approach to derive minimizer schemes. These schemes not only demonstrate provably lower density compared to random minimizers and other existing schemes but are also fast to compute (even in streaming fashion), do not require any auxiliary space, and are easy to analyze. Notably, a specific instantiation of the framework gives a scheme, the *mod-minimizer*, that achieves optimal density when *k* → ∞. Although Marçais et al. [11] were the first to describe a method that achieves optimal density when *k* → ∞, the proof of optimality of the mod-minimizer is simple and does not rely on complex machinery as previous approaches. (As a side contribution, we review such previous approaches using a consistent notation and we simplify their exposition.) The mod-minimizer has lower density than the method by Marçais et al. for practical values of *k* and *w* and converges to 1*/w* faster.

Our theoretical analysis matches empirical performance on both synthetic and real-world strings: the mod-minimizer significantly outperforms the random minimizer and other state-of-the-art methods like closed syncmers [4] and miniception [27] in practice, given the appropriate choice of parameters. We integrated the mod-minimizer into SSHash [18, 19], a *k*-mer dictionary based on minimizers, resulting in a substantial reduction in space usage (14–15%) while maintaining its fast query times. Both C++ and Rust implementations of the proposed schemes are publicly available on GitHub.

## 2 Notation, preliminary definitions, and problem statement

We report here the notation used throughout the paper, along with some useful definitions.

### Notation

Let [*n*]:= {0, …, *n* −1}, for any *n* ∈ ℕ = {1, 2, 3, …}. We fix an alphabet Σ = [*σ*] of size *σ* = 2^*O*(1)^. Let *S* ∈ Σ^∗^ be a string. We refer to *S*[*i*..*j*) as its sub-string of length *j* − *i* starting at index *i* and ending at index *j* (excluded). When *j* − *i* = *k* for some *k* ≥ 1, we call *S*[*i*..*j*) a *k*-mer of *S*. In the following, let *w >* 0 be an integer, so that any string of length *ℓ* = *w* + *k* − 1 defines a *window W* of *w* consecutive *k*-mers. We refer to *w* as the *size* of the window. (Whenever we use the term “window” in the following, we implicitly assume that to be relative to an hypothetical long string *S*.) It follows that each *k*-mer in *W* can be uniquely identified with an integer in [*w*], corresponding to its starting position in *W*. We say that two windows *W* and *W* ^*′*^ are *consecutive* when *W* [1..*ℓ*) = *W* ^*′*^[0..*ℓ* − 1).

We write *a* mod *m* for the remainder of *a* after division by *m* and *a* ≡ *b* (mod *m*) to say that *a* and *b* have the same remainder modulo *m*.

#### Definition 1

(Order). *An order* 𝒪_*k*_ *on k-mers is a function* 𝒪_*k*_: Σ^*k*^ → ℝ, *such that* 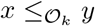 *if and only if* 𝒪_*k*_(*x*) ≤ 𝒪_*k*_(*y*).

We do not necessarily require 𝒪_*k*_ to be *random*, although practitioners often use a (pseudo-)random hash function *h*: Σ^*k*^→ [*U*] to define the order, where [*U*] is a sufficiently large range, like *U* = 2^128^. We therefore make the standard assumption [1, 21] that *h* is drawn from a family of fully random hash functions that can be evaluated in *O*(1) on a machine word.

### Hashing *k*-mers

Since *σ* = 2^*O*(1)^, it follows that any *k*-mer *x* ∈ Σ^*k*^ fits in *O*(*k*) words and *h*(*x*) is computed in *O*(*k*) time. Furthermore, using a rolling hash function [13], we can compute *w* hashes for the *w* consecutive *k*-mers in a window in *O*(*w* + *k* − 1) rather than the naïve *O*(*wk*). We implicitly assume this linear bound hereafter when discussing the complexities of the algorithms.

### Sampling functions

We now define a hierarchy of *sampling* functions.

#### Definition 2

(Local scheme). *A* local scheme *is a function f*: Σ^*w*+*k*−1^ → [*w*] *that, given a window W*, *selects the k-mer starting at position f*(*W*) *in W*. *That is, the sampled k-mer is W* [*f*(*W*)..*f* (*W*) + *k*).

#### Definition 3

(Forward scheme). *A local scheme f is a* forward scheme *when for any two consecutive windows W and W* ^*′*^ *it holds that f*(*W*) ≤ *f*(*W* ^*′*^) + 1.

Forward schemes have the property that as the window *W* slides through an input string *S*, the position in *S* of the selected *k*-mer never decreases. This is, for example, the case for minimizers as defined below.

#### Definition 4

(Minimizer scheme). *A* minimizer scheme *is defined by an order* 𝒪_*k*_ *on k-mers and selects the leftmost minimal k-mer in a window W*, *which is called the* minimizer:

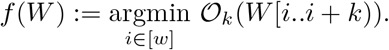

#### Definition 5

(Random minimizer). *The* random minimizer *is the minimizer scheme when* 𝒪_*k*_ *is a random order*.

The performance metric we focus on in this work is the *density* of a scheme.

#### Definition 6

(Density). *Given a string S of length n, let W*_*i*_:= *S*[*i*..*i* +*ℓ*) *for i* ∈ [*n* −*ℓ* + 1]. *A sampling function f selects the k-mers starting at positions* {*i* + *f*(*W*_*i*_) | *i* ∈ [*n* − *ℓ* + 1]}. *The* particular density *of on is* |{*i + f(W*_*i*_) | *i* ∈ *[n − ℓ + 1]*} /*(n − k + 1). The density d*(*f*) *of f is defined as the expected particular density on a string S consisting of i*.*i*.*d. random characters of* Σ *in the limit where n* → ∞.

We remark that Marçais et al. [11] give a definition of density in terms of particular density on *circular* strings (which we do not define here for brevity). For *n* → ∞, their definition and Definition 6 are equivalent. Hence, we use finite but sufficiently-long random strings to approximate the density in this work.

Random minimizers have been the most popular choice by practitioners because they are simple to implement and offer overall good performance in terms of sampling speed and density. Let *R* be a minimizer scheme where 𝒪_*k*_ is a random *total* order. Then, the density of *R* is *d*(*R*) ≈ 2/(*w* + 1) [26]^1^. However, there are schemes with lower density. Let the sets of all local schemes, all forward schemes, and all minimizer schemes be denoted by ℒ, ℱ, and ℳ respectively. All minimizer schemes are forward [11], and by definition all forward schemes are local, so that ℳ ⊆ ℱ ⊆ ℒ. We write *d*(ℒ), *d*(ℱ), and *d*(ℳ) for the best possible density of a local, forward, and minimizer scheme respectively. We have

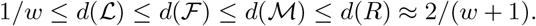

With these initial remarks in mind, we can precisely state the problem under study here.

#### Problem 1

(`Pure sampling function problem`). *Given integers w* ≥ 2 *and k* ≥ 1, *implement a function f*: Σ^*w*+*k*−1^ → [*w*] *in O*(1) *space with as low density as possible*.

A few considerations about Problem 1 are in order.

▪ By definition, any function solving Problem 1 automatically satisfies the window guarantee that at least one *k*-mer is sampled from every *w* consecutive *k*-mers from a string.
▪ We require *w* ≥ 2, since for *w* = 1 the density of any scheme is just trivially 1.
▪ It is desirable that the evaluation time of *f*, referred to as *sampling time* in the following, is proportional to the number of characters in the window or faster, i.e., *O*(*w* + *k* − 1).

## 3 Review of previous methods

In this section we review previous methods, in chronological order of proposal. Our focus is on the algorithmic description of the methods; for a survey about applications of minimizers, refer to [29]. In particular, we are interested in methods that solve Problem 1, i.e., that (1) can be implemented using constant space and (2) have a window guarantee. These methods are: the random minimizer [23, 26], the “rotational” minimizer [11], the miniception [27], the closed syncmer [4], and the minimum decycling set based minimizers [17].

In the following, we describe these approaches and give concise pseudocode to aid concrete implementations. Although the pseudocode illustrates how the pure function *f*: Σ^*w*+*k*−1^ → [*w*] from Problem 1 can be implemented, we point out that each of the methods can be implemented in a *streaming* fashion as well, that is, they can be implemented so that it takes *O*(*m*) amortized time to compute *f* for *m* consecutive windows. (Our concrete implementations support both stateless and streaming sampling.)

### Algorithm 1

Pseudocode for the random minimizer algorithm.

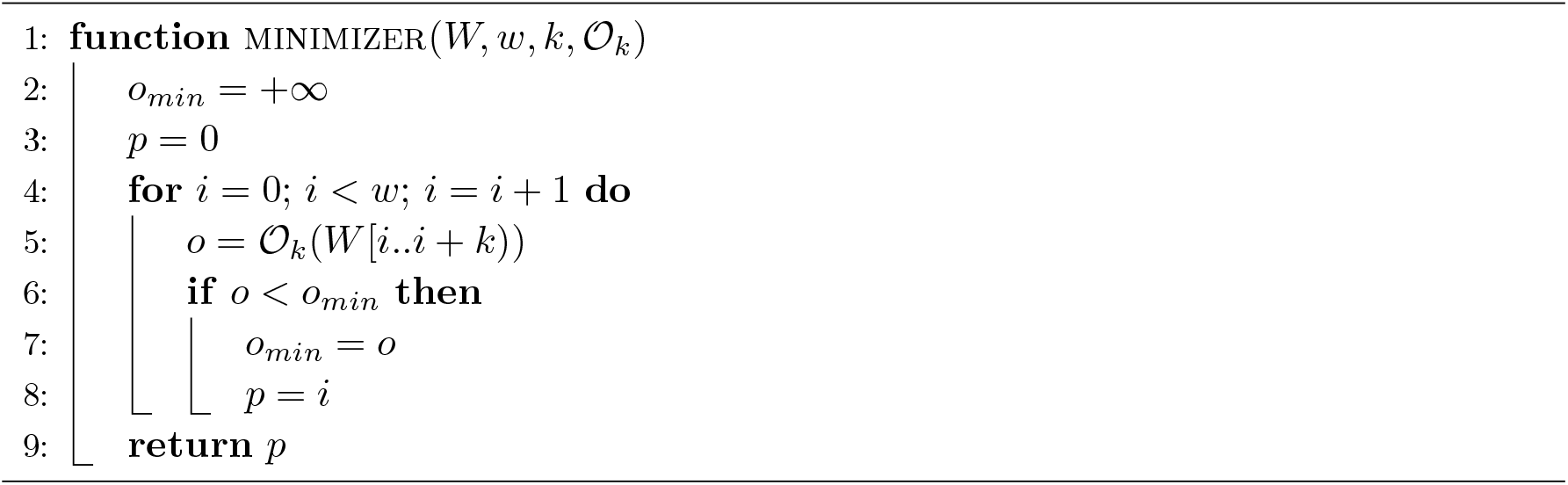

### Random minimizer

As per Definition 4, the minimizer of a window is the smallest *k*-mer according to some order 𝒪_*k*_. Computing a minimizer thus takes *O*(*w* + *k* − 1) time and the relevant pseudocode is given in Algorithm 1. As already mentioned, the minimizer has density ≈ 2/(*w* + 1) when 𝒪_*k*_ is a random total order [26].

### Rotational minimizer

Marçais et al. [11] present a scheme based on a universal hitting set (UHS) that approaches density 1*/w* when *w* is fixed and *k* → ∞. A UHS is a set of *k*-mers such that *every* string of length *ℓ* = *w* + *k* − 1 contains at least one *k*-mer in the UHS [15]. Hence, in the large-*k* limit, the hierarchy described in Section 2 collapses and minimizers are as good as forward and local schemes. Assuming that *w* divides *k*, for each *j* ∈ [*w*] this scheme considers the sum *ψ*_*j*_ of characters (with values in [*σ*]) in positions *j* mod *w* in each *k*-mer in the window. A *k*-mer is part of the UHS when *ψ*_0_ + *σ* ≥ max_*j*∈ [*w*]_ *ψ*_*j*_ holds true. In the minimizer scheme, we break ties by falling back to a random minimizer. We call this scheme the “rotational” minimizer^2^, inspired by the geometrical rotation of the (*ψ*_0_, …, *ψ*_*w*−1_) embedding. The complexity of Algorithm 2 is *O*(*wk*). The pseudocode in Algorithm 2 is faithful to the description from the original paper [11]. Algorithm 7 in the Appendix shows our own modified version which gives better results than the original in practice, by simply preferring *k*-mers with large ψ_0_.

### Miniception

The term *miniception* stands for “minimizer inception” and the method is based on the idea of using two minimizers with different parameters [27]. Details are given in Algorithm 3. For a given parameter *t* ≤ *k*, the miniception samples the smallest *k*-mer such that the position of its smallest contained *t*-mer is either 0 or *k* − *t*. When *t < k* − *w*, such *charged k*-mers may not exist and the smallest *k*-mer is sampled. In our implementation and experiments we always use the recommended *t* = max(*k − w*, 3). When *t* = *k* − *w* + 1, the density achieved by the miniception can be upper bounded by 1.67*/w* + *o*(1*/w*) when 𝒪_*t*_ and 𝒪_*k*_ are random total orders ([27], Theorem 7). In practice, *t* = *k* − *w* is used. The identification of the smallest *t*-mer (the call to minimizer(*W* [*i*..*i* + *k*), *w*^*′*^ + 1, *t*, 𝒪_*t*_) in Algorithm 3) can be done in streaming fashion, so that the complexity of miniception is linear in the length of the window, i.e., *O*(*w* + *k* − 1), and not quadratic.

### Closed syncmer

For a given parameter *t* ≤ *k*, a *k*-mer is a *closed syncmer* [4] if its smallest *t*-mer is in position 0 or *k* − *t*, corresponding to the charged *k*-mers of miniception. Note that this definition is “context free”, in that it does not depend on surrounding *k*-mers. The first *k*-mer in a window that is a closed syncmer is sampled, as illustrated in Algorithm 4. As already noted, the identification of the smallest *t*-mer in each of the *k*-mers in a window can be implemented in linear time, so that the complexity of Algorithm 4 is *O*(*w* + *k* − 1). Closed syncmers satisfy a window guarantee *w* when *t* ≥ *k* − *w*. The closed syncmer has a density of 2*/*(*k* − *t* + 1) (assuming a random total order on *t*-mers), which is the same as that of a random minimizer when *t* = *k* − *w*. Syncmers were indeed designed to improve the *conservation* metric rather than density compared to minimizers (see the original paper by Edgar [4] for details). Note that closed syncmers differ from miniception in that they consider *all* charged *k*-mers, as opposed to considering only those that have a small order 𝒪_*k*_.

#### Algorithm 2

Pseudocode for the rotational minimizer algorithm.

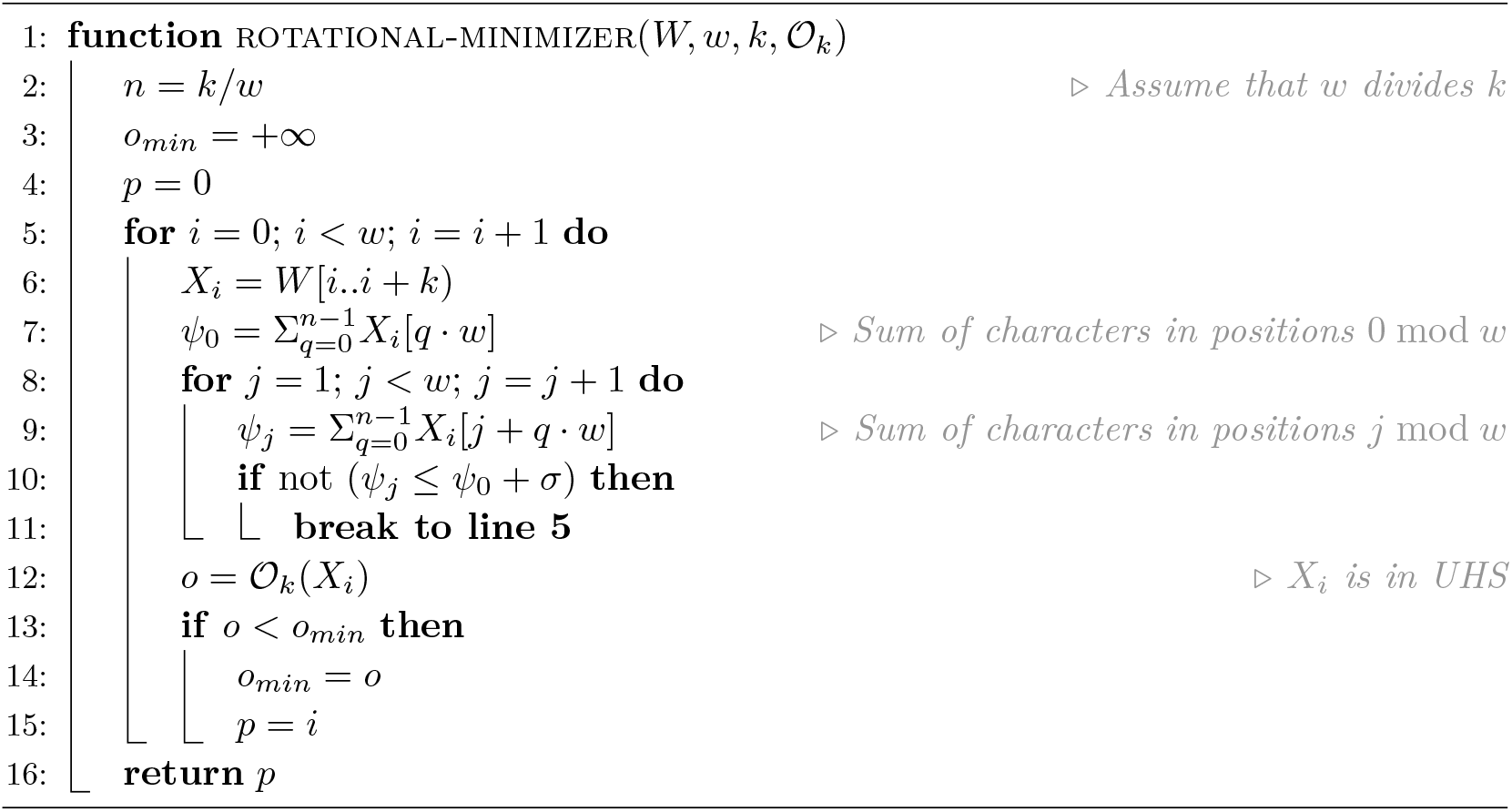

### Minimum decycling set

A *decycling set* is a set of *k*-mers such that any infinitely long string contains a *k*-mer in the set. Mykkeltveit [14] shows how to construct a minimum decycling set (MDS) using an embedding of *k*-mers into the complex plane. In particular, each *k*-mer *X* is mapped to a complex point *x* resulting from the weighted sum of the *k*-th complex roots of unity (line 7 of Algorithm 5). In this way, a cyclic rotation of *X* to the left, i.e., left-rotation(*X*) = *X*[1..*k*) · *X*[0], corresponds to a rotation in *clockwise* direction of the complex number *x* by an angle of 2*π/k*. The MDS 𝒟_*k*_ constructed by Mykkeltveit consists of those *k*-mers whose embedding corresponds to the *first clockwise rotation having positive imaginary part*, i.e., such that *π* − 2*π/k* ≤ arg(*x*) < π.

Pellow et al. [17] use 𝒟_*k*_ to construct a minimizer order as follows^3^: all *k*-mers in 𝒟_*k*_ are smaller than those in Σ^*k*^ \ 𝒟_*k*_; within 𝒟_*k*_ and Σ^*k*^ \ 𝒟_*k*_, all *k*-mers are relatively ordered by a given 𝒪_*k*_. They also define a *double* decycling-set-based order using the *symmetric* MDS 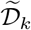, the set of *k*-mers for which −2*π/k* ≤ arg(*x*) *<* 0, and then break ties in favour of 𝒟_*k*_ first, then 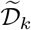, and lastly 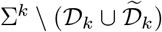. See Algorithm 5, with complexity *O*(*wk*).

### Other methods

Several related sampling algorithms have been proposed that do not solve Problem 1. Universal hitting sets that explicitly list *k*-mers [16, 6] consume more than *O*(1) space. Open syncmers [4] and minmers [8] do not consistenly have a small window guarantee. Bidirectional anchors sample *positions* rather than *k-mers* [10, 9]. Lastly, sequence-specific schemes [28] can achieve very low density but consume more than *O*(1) space.

#### Algorithm 3

Pseudocode for the miniception algorithm.

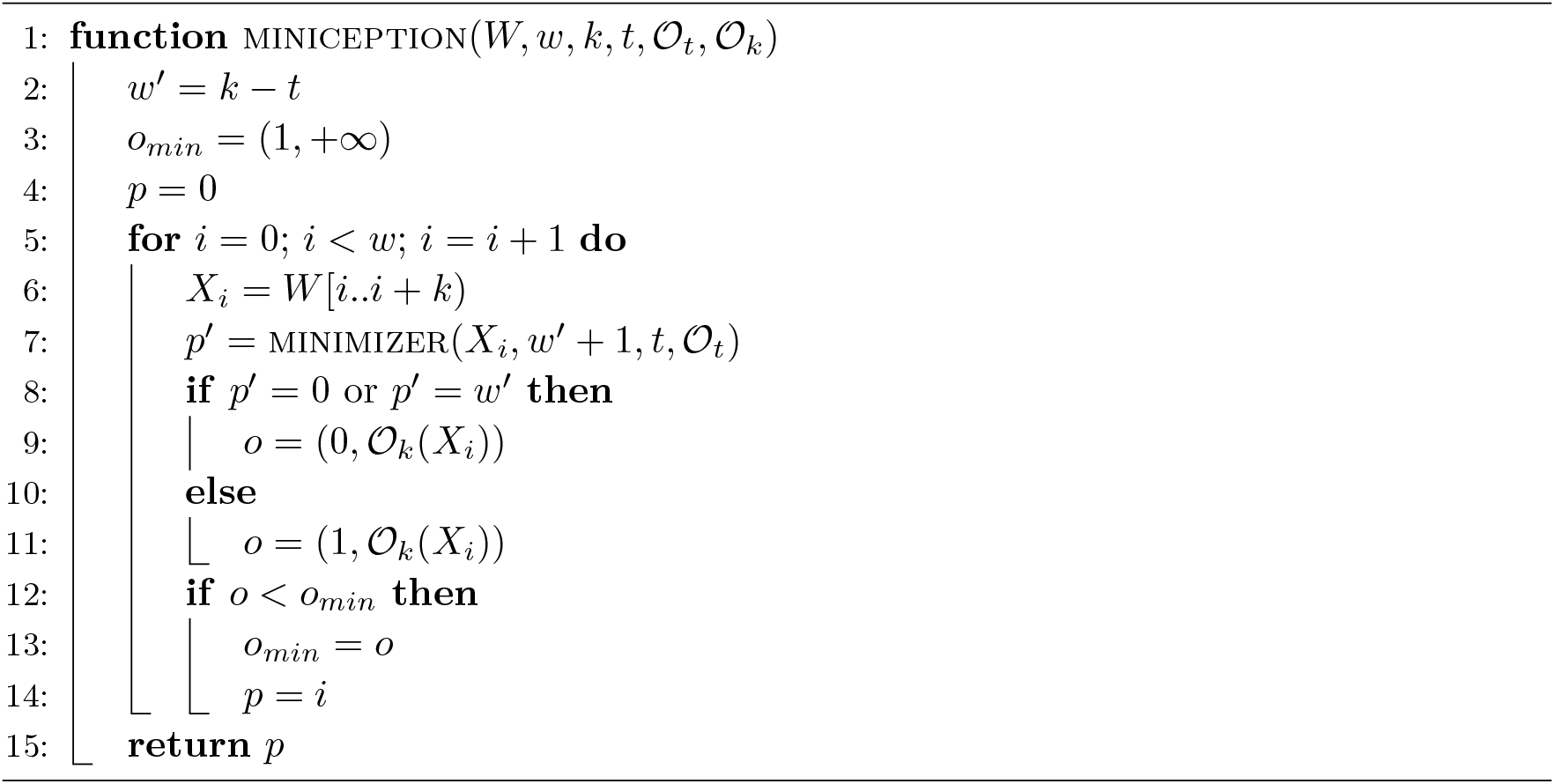

#### Algorithm 4

Pseudocode for the closed syncmer algorithm.

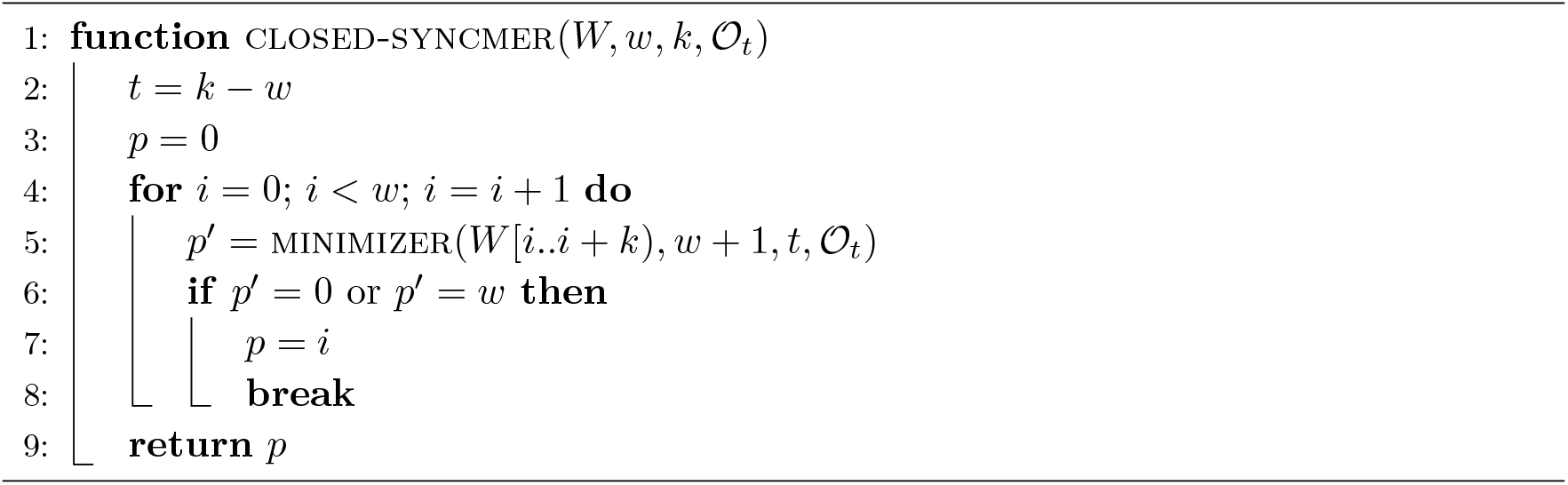

## 4 New minimizer schemes using modulo sampling

This section presents our main contribution, the *mod-minimizer*, that has provably lower density than the random minimizer and achieves optimal density 1*/w* for *large k*. To be precise, when we say “large *k*” in the following, we mean for *w* fixed and *k* → ∞.

### 4.1 Modulo sampling

The mod-minimizer is obtained as a special case of a more general two-step sampling algorithm, that we name *modulo sampling* (or just *mod-sampling* for brevity).

#### Definition 7

(Mod-sampling). *Let W be a window containing w k-mers. Let* 1 ≤ *t* ≤ *k be a given parameter, and* 𝒪_*t*_ *be an order on t-mers. Let S be the set of starting positions of the minimal t-mers in W*. *Then, sample the k-mer starting at position* min{*i* mod *w* | *i* ∈ *S*}.

The pseudocode of mod-sampling is given in Algorithm 6. It is straightforward to see that its complexity is *O*(*w* + *k* − 1). The mod-sampling algorithm defines a two-step approach where the first step is to find the positions of the minimal *t*-mers, followed by the identification of the actual sampled *k*-mer via modulo.

#### Algorithm 5

Pseudocode for the minimum decycling set algorithm.

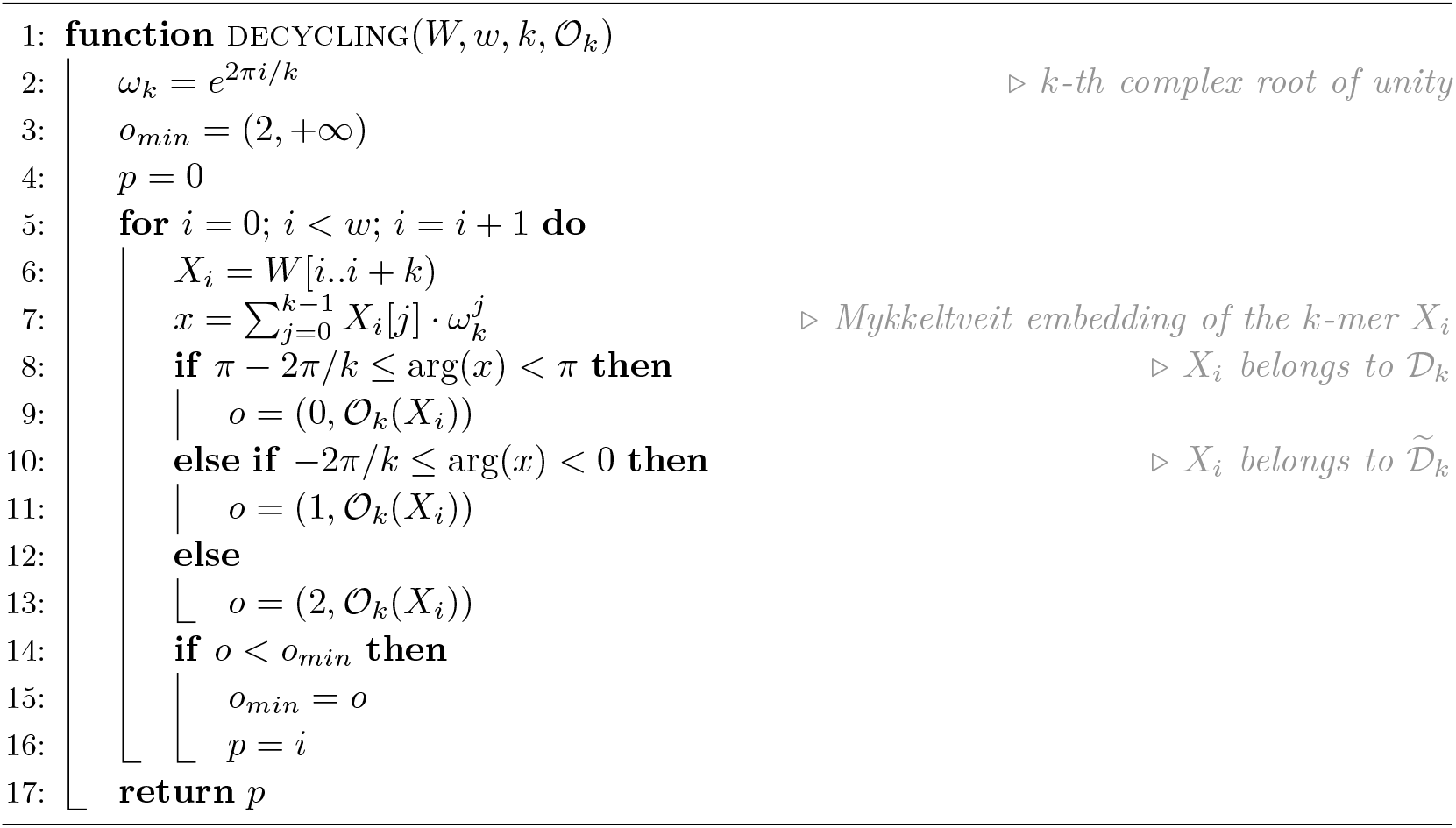

#### Algorithm 6

The pseudocode for the mod-sampling algorithm (Definition 7).

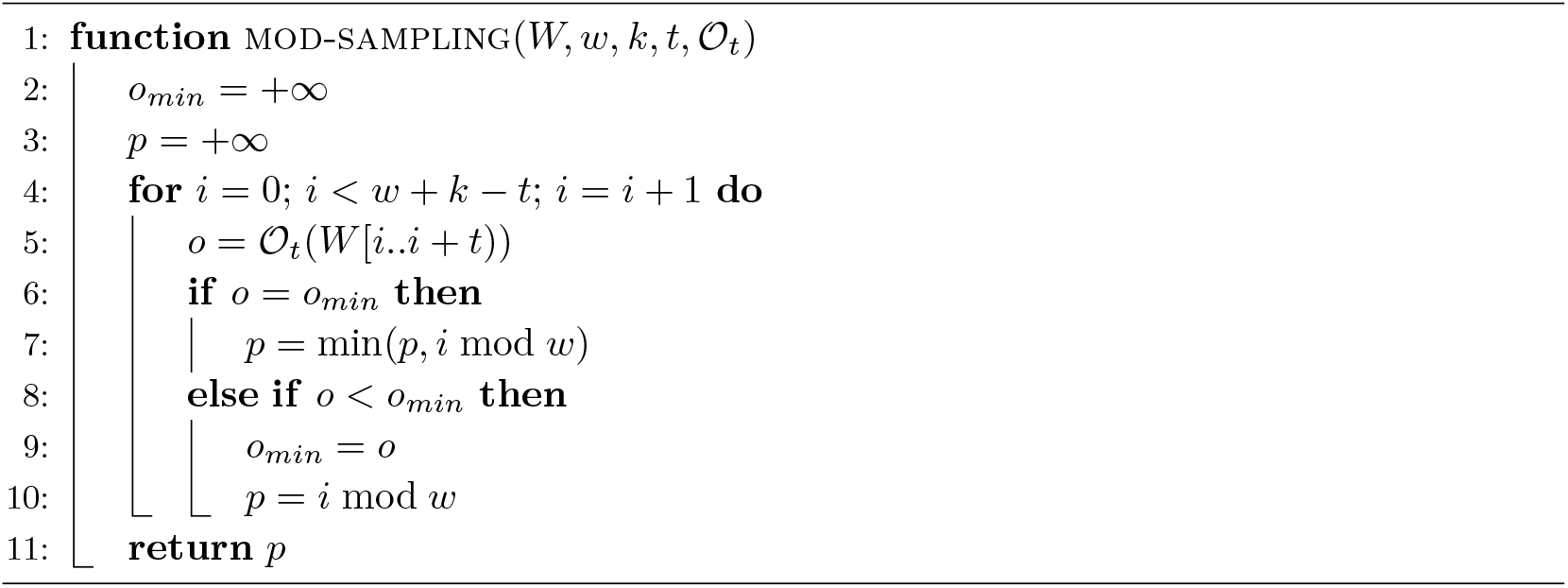

From now on, we assume that the order on *t*-mers 𝒪_*t*_ is *random* (and, as already noted in Section 2, in practice we use a pseudo-random hash function with a large output range).

#### Why mod-sampling intuitively works well for large *k*

Before digging into technical details, we first give some intuition. Let *w* be fixed and assume that *k* is large compared to *w*. For now we will assume that *t* is a small constant at most *w*, and hence also small compared to *k*.

Assume, for simplicity, that there is one occurrence of the minimal *t*-mer in the window. As long as the minimal *t*-mer in the window remains the same, mod-sampling will either sample the same *k*-mer for consecutive windows, or shift the position of the sampled *k*-mer by exactly *w* steps, as can be seen in Figure 1. The window guarantee requires us to sample at least one every *w* positions, so locally, this is the best we can do.

**Figure 1.**
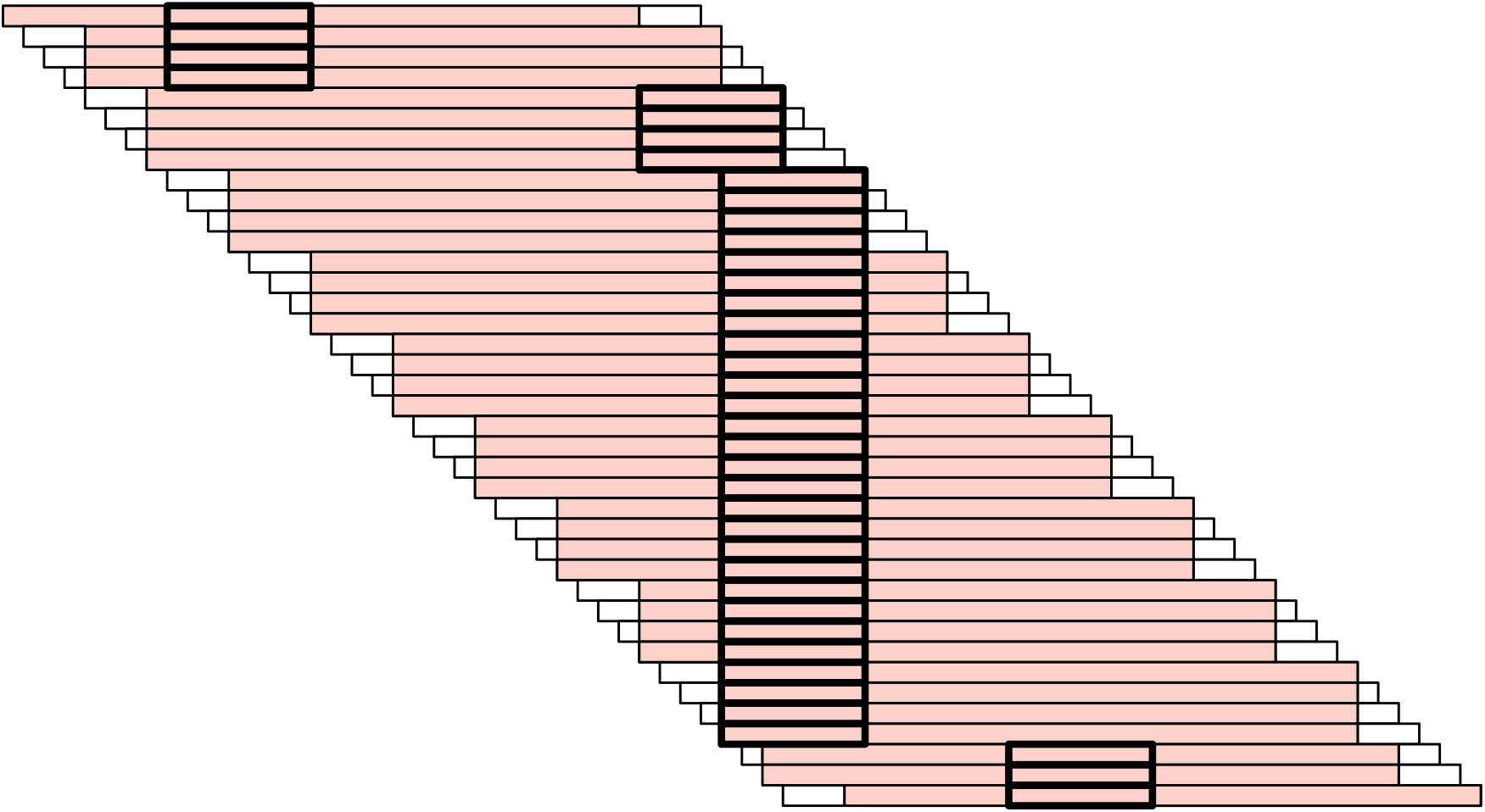
An illustration of mod-sampling for *k* = 31, *w* = 4, and *t* = 7. Rows indicate consecutive windows. The thick outlined boxes mark the minimal *t*-mer in each window and the regions highlighted in red indicate the sampled *k*-mer. When the minimal *t*-mer is preserved over many windows, which is likely when *k* is large compared to *t* and *w*, this *t*-mer will serve as an “anchor”, causing *the same k*-mer to be sampled in blocks of *w* consecutive windows, effectively sampling one every *w k*-mers.

The sampled *k*-mer can also change when the minimal *t*-mer changes. Thus, we would like the minimal *t*-mer to change *as infrequently as possible*. Since *t* is small compared to *k*, the number of *t*-mers in each window of length *ℓ* = *w* + *k* − 1 is *ℓ* − *t* + 1 = *w* + (*k* − *t*), which is much larger than the number of *k*-mers, *w*. This means that minimal *t*-mers are preserved over longer distances compared to minimal *k*-mers (i.e., the expected distance is (*w* + (*k* − *t*) + 1)*/*2 rather than (*w* + 1)/2, effectively improving over random minimizers). Now, as we take *k* → ∞, we see that the length over which minimal *t*-mers are preserved also goes to ∞. This means that in the limit, the density of the mod-sampling scheme is fully determined by jumps of size *w*, and hence the density converges to 1*/w*.

One issue is that as the windows get infinitely long, they will exceed length σ^*t*^ and contain duplicate *t*-mers. To mitigate this effect, we let *t* grow as Ω(log(*w* + *k* − 1)), which is still small enough to ensure that the density of minimal *t*-mers converges to 0 (for details, see Lemma 9).

Depending on the choice of *t*, mod-sampling can yield *non-forward* schemes as proven in Lemma 8. Consider the example in the bottom two rows of Figure 2a: *w* = 4, *k* = 6, *t* = 5, and let *W* and *W* ^*′*^ be two consecutive windows. Window *W* may have the minimal *t*-mer in position 3, giving *f*(*W*) = 3 mod 4 = 3, and *W* ^*′*^ can introduce a new smaller *t*-mer in its last position *w* + *k* − *t* − 1 = 4, so that *f*(*W* ^*′*^) = 4 mod 4 = 0 *< f*(*W*) − 1 = 2. We see that *W* ^*′*^ samples a *k*-mer to the left of the one sampled by *W* and we say that *W* ^*′*^ introduces a “backward jump”.

**Figure 2.**
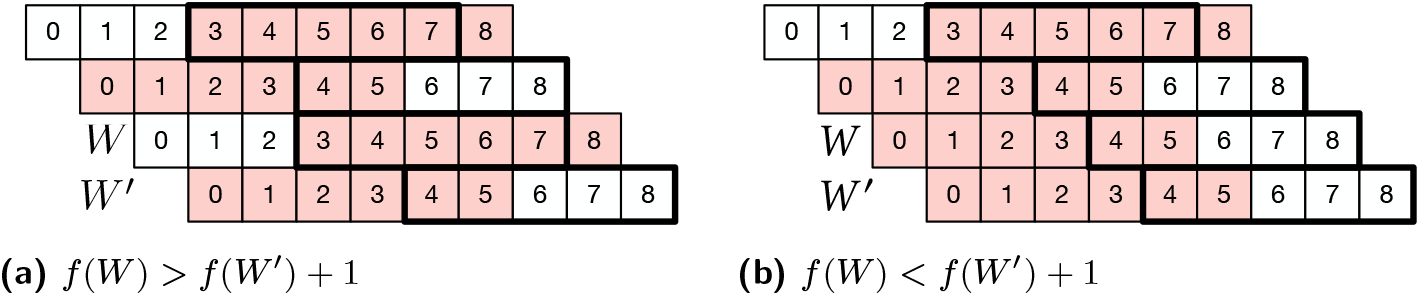
Examples of doubly sampled *k*-mers due to backward jumps for *w* = 4, *k* = 6, and *t* = 5, where the thicker stroke marks the minimal *t*-mer and red boxes indicate the sampled *k*-mer from each window. In both these examples, backward jumps cause the *k*-mer *W* ^*′*^[*f* (*W* ^*′*^)..*f* (*W* ^*′*^) + *k*) be sampled twice.

##### Lemma 8.

*Mod-sampling is forward if and only if t* ≡ *k* (mod *w*) *or t* ≡ *k* + 1 (mod *w*).

**Proof**. Consider two consecutive windows *W* and *W* ^*′*^. Let *i* be the position of the minimal *t*-mer in window *W* and *i*^*′*^ that of the minimal *t*-mer in *W* ^*′*^. Mod-sampling is forward when (*i* mod *w*) − 1 ≤ (*i*^*′*^ mod *w*) holds for all *i* and *i*^*′*^. Given that the two windows are consecutive, this trivially holds when *i* = 0 and when *i*^*′*^ = *i* − 1. Thus, the only position *i*^*′*^ that could violate the forwardness condition is when *W* ^*′*^ introduces a new minimal *t*-mer at position *i*^*′*^ = *w* +*k* −*t*− 1. In this case, we have *i*^*′*^ mod *w* = (*w* +*k* −*t*− 1) mod *w* = (*k* −*t*− 1) mod *w*. The rightmost possible position of the sampled *k*-mer in *W* is *i* mod *w* = *w* − 1. Hence, if the scheme is forward, then it must hold that (*w* − 1) − 1 = *w* − 2 ≤ (*k* − *t* − 1) mod *w*. Vice versa, if *w* − 2 ≤ (*k* − *t* − 1) mod *w* always holds true, then the scheme is forward.

Now, note that (*k* − *t* − 1) mod *w* ≥ *w* − 2 ⇔ *qw* − 2 ≤ *k* − *t* − 1 *< qw* ⇔ *k* − *qw* ≤ *t* ≤ *k* − *qw* + 1 for some 0 ≤ *q* ≤ ⌊*k/w*⌋. In conclusion, the scheme is forward if and only if *t* = *k* − *qw* or *t* = *k* − *qw* + 1, i.e., when *t* ≡ *k* (mod *w*) or *t* ≡ *k* + 1 (mod *w*).

The next theorem gives the expression for the density of a random scheme yielded by the mod-sampling algorithm, but first we give a useful lemma based on Lemma 9 of [27]. The proof is in Appendix A.1.

##### Lemma 9.

*For any* ϵ > 0, *if t >* (3 + *ϵ*) log_*σ*_(*ℓ*), *the probability that a random window of ℓ* − *t* + 1 *t-mers contains two identical t-mers is o*(1*/ℓ*). *Given that ℓ* = *w* + *k* − 1, *o*(1/*ℓ*) → 0 *for large k*.

##### Theorem 10.

*If t is chosen as in Lemma 9 and* 𝒪_*t*_ *is a random total order, the density of the mod-sampling scheme is at most*

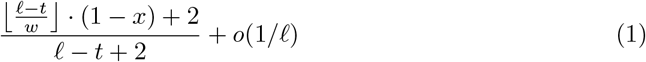

*where*

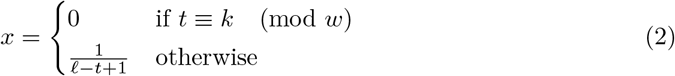

*and ℓ* = *w* + *k* − 1.

**Proof**. Consider two consecutive windows *W* and *W* ^*′*^ of length *ℓ* of a uniform random string. For forward schemes, the density can then be computed as the probability that a different *k*-mer is sampled from *W* ^*′*^ than from *W*, i.e., ℙ[*f*(*W*) ≠ *f*(*W* ^*′*^)+1]. However, for non-forward schemes, we cannot be sure that the *k*-mer *W* ^*′*^[*f*(*W* ^*′*^)..*f* (*W* ^*′*^) +*k*) was not already sampled in any of the previous *w* − 2 windows because of possible backward jumps. Figure 2 illustrates some examples. Therefore, in general, ℙ[*f*(*W*) ≠ *f*(*W* ^*′*^) + 1] only provides an *upper bound* to the density of mod-sampling. For brevity, we write *Z* for the indicator random variable that a different *k*-mer is sampled from *W* ^*′*^ than from *W*. The density is then at most 𝔼[*Z*] = ℙ[*f*(*W*) ≠ *f*(*W* ^*′*^) + 1].

Let *U* be the event that the minimal *t* -mer according to an order 𝒪_*t*_ in an (*ℓ* + 1)-mer is unique. By Lemma 9, ℙ [*Ū]* = *o*(1/*ℓ*).

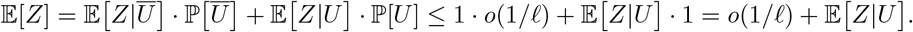

In the remainder we assume that the minimum is unique to bound 𝔼 [*Z* |*U]*.

We write *i* and *i*^*′*^ for the position of the minimal *t*-mer in *W* and *W* ^*′*^ respectively and, similarly, we let *p* = *f*(*W*) and *p*^*′*^ = *f*(*W* ^*′*^). Let ≤ 0 ≤ *y ℓ* − *t* + 1 be the position of the minimal *t*-mer in the union of the two windows. We distinguish between three cases (refer to Figure 3).

**Figure 3.**
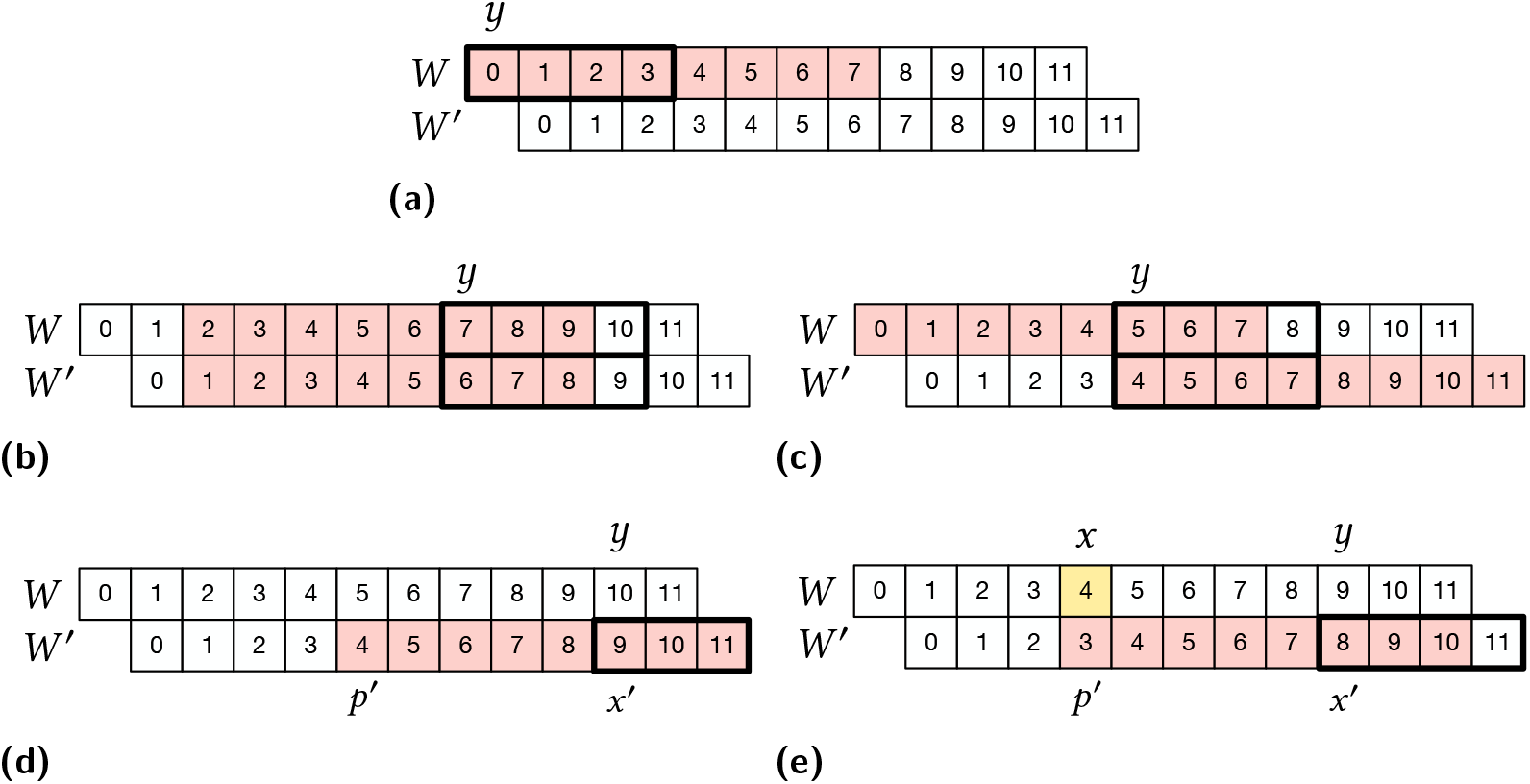
The different cases used in the proof of Theorem 10, depending on the position *y* of the minimal *t*-mer in two consecutive windows, *W* and *W* ^*′*^. All examples are for *k* = 8, *w* = 5, and *t* = 4 except for case **(d)** where we use *t* = 3.

▪ Case *y* = 0. In this case the first window has its minimal *t*-mer at *i* = 0 and hence *p* = 0, and the two windows must necessarily sample different *k*-mers (Figure 3a).
▪ When 0 *< y < ℓ* − *t* + 1, the minimal *t*-mer is shared between the two windows. We therefore have *p* = *y* mod *w* and *p*^*′*^ + 1 = ((*y* − 1) mod *w*) + 1. When *y* is not a multiple of *w*, we have *p* = *p*^*′*^ + 1 and the same *k*-mer is sampled (Figure 3b). When *y* is a multiple of *w*, i.e., *y* = *w*, 2*w*, 3*w*, …, ⌊(*ℓ* − *t*)*/w*⌋*w, W* ^*′*^ samples a new *k*-mer exactly *w* to the right of the *k*-mer sampled by *W* (Figure 3c). There are ⌊(*ℓ* − *t*)*/w*⌋ such positions *y*.
▪ When *y* = *ℓ* − *t* + 1, the minimal *t*-mer in *W* ^*′*^ is not present in *W*. If *p*^*′*^ = *i*^*′*^ mod *w* = (*y* − 1) mod *w* = (*ℓ* − *t*) mod *w* equals *w* − 1, then the two windows must necessarily sample different *k*-mers (Figure 3d). Otherwise, i.e., if (*ℓ* − *t*) mod *w < w* − 1, the two windows sample the same *k*-mer when ((*ℓ* − *t*) mod *w*) + 1 = *p*^*′*^ + 1 = *p* = *i* mod *w*. The number of such positions *i* is ⌊(*ℓ* − *t*)*/w*⌋ (one such position is, for example, the one depicted in yellow in Figure 3e). Because 𝒪_*t*_ is random, *i* takes each position with uniform probability 1*/*(*ℓ* − *t* + 1). Hence, with probability 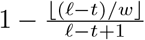 the two windows sample different *k*-mers in this case.

Summing up, as the minimal *t*-mer can be in any position *y* with probability 1*/*(*ℓ* − *t* + 2) because 𝒪_*t*_ is random, we conclude that 𝔼[*Z*] is at most

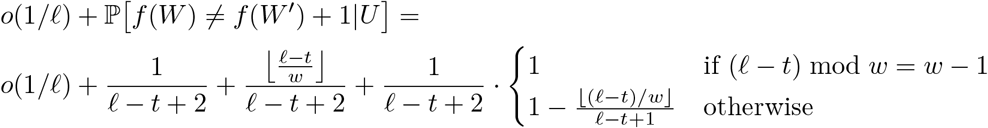

and, by simplifying, the claim follows.

The following corollary shows that mod-sampling achieves optimal density for large *k*.

##### Corollary 11.

*If t is chosen as in Lemma 9, the mod-sampling scheme has optimal density for large k, i*.*e*., *the density of mod-sampling converges to* 1*/w as w is fixed and k* → ∞.

**Proof**. First note that density of Theorem 10 can be bounded from above by always taking *x* = 0. Since *k* → ∞ and *t* = *o*(*ℓ*), we also have (*ℓ* − *t*) → ∞, so we obtain

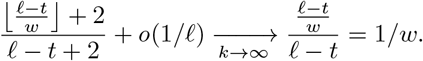

The next lemma, proven in Appendix A.4, describes the shape of the density function in Formula 1 when *ℓ* and *w* are fixed, and *r* ≤ *t* ≤ *k*. Although mod-sampling is well defined for any 1 ≤ *t* ≤ *k*, in practice we restrict our attention to *t* ≥ *r* for some lower bound *r* to mitigate the effect of duplicate *t*-mers in a window.

##### Lemma 12.

*The density function in Formula 1 has a “sawtooth” shape: for any integer* 1 ≤ *r* ≤ *k, it achieves its minimum on r* ≤ *t* ≤ *k either in t* = *r or in t* = *r* + ((*k* −*r*) mod *w*).

Consider Figure 4 for some concrete examples. As clearly visible from those examples, our analysis is tight for forward schemes and, for those cases that are not forward, the computed density provides an upper bound that is close to the measured density. Only for very small *t*, like 2 and 3, duplicate *t*-mers sometimes cause the measured density to exceed the computed density, but these cases are not relevant in practice. The gap between computed and measured density is most visible for *t* = *k* − 1, and is never more than 1.7% in the given examples. Furthermore, Lemma 8 together with Lemma 12 imply that the forward schemes given by mod-sampling minimize the density in Formula 1 and, hence, are more powerful than its local schemes.

**Figure 4.**
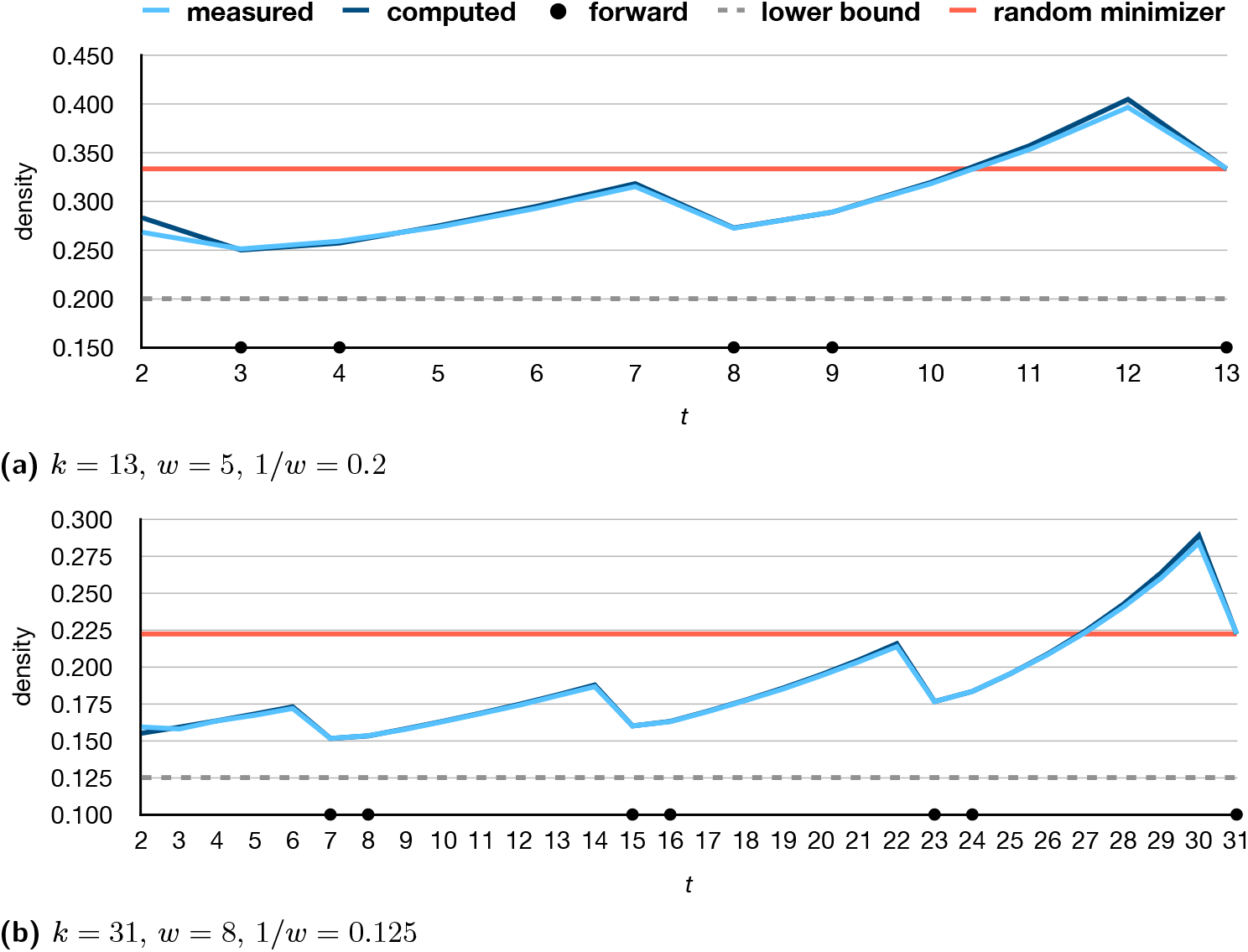
Two examples, for different *k* and *w*, of the density of mod-sampling. In particular, the plots show the measured density by varying *t* (in light blue), calculated over a random string of one million characters with *σ* = 4 in comparison to the density computed using Formula 1 (in dark blue), neglecting lower order terms. For *r* ≤ *t* ≤ *k*, the density is minimum at *t* = *k* mod *w*. (Here, we use *r* = 2 to avoid degenerate cases.) Black dots indicates the values of *t* for which the scheme is forward. As a reference point, the red straight line corresponds to 2*/*(*w* + 1), the density of a random minimizer scheme.

However, we remark that *t* = *k* mod *w* could be too small, for example when *w* = 8 and *k* ∈ {32, 33, 34}. In this case one should then pick *t* = (*k* mod *w*) + *w* to mitigate the effect of duplicate *t*-mers in a window^4^. In the light of this analysis, we focus hereafter on the choice *t* ≡ *k* (mod *w*).

### 4.2 The mod-minimizer

Now that we have presented the mod-sampling algorithm and analyzed its performance, in this section we fix some concrete choices of *t* such that *t* ≡ *k* (mod *w*). For these choices, the following lemma shows that we obtain *minimizer* schemes.

#### Lemma 13.

*The mod-sampling algorithm yields a minimizer scheme if t* ≡ *k* (mod *w*).

**Proof**. Our proof strategy explicitly defines an order 𝒪_*k*_ and shows that mod-sampling with *t* ≡ *k* (mod *w*) corresponds to a minimizer scheme using 𝒪_*k*_, i.e., the *k*-mer sampled by mod-sampling is the leftmost minimal *k*-mer according to 𝒪_*k*_.

Let 𝒪_*t*_ be the order on *t*-mers used by mod-sampling. Define the order 𝒪_*k*_(*X*) of the *k*-mer *X* as the order of its minimal *t*-mer, chosen among the *t*-mers occurring in positions that are a multiple of *w*:

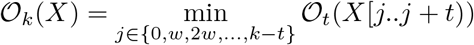

where *k* − *t* is indeed a multiple of *w* since *t* ≡ *k* (mod *w*).

Now consider a window *W* of consecutive *k*-mers *X*_0_, …, *X*_*w*−1_. Since each *k*-mer starts at a different position in *W*, 𝒪_*k*_(*X*_*p*_) considers different sets of positions relative to *W* than 𝒪_*k*_(*X*_*q*_) for all *p* ≠ *q* (but *t*-mers starting at different positions in *W* could have the same order, i.e., the minimal *t*-mer of *X*_*p*_ could have the same order as that of *X*_*q*_). These sets form a perfect, non-overlapping partition of every single *t*-mer in the entire window. Every absolute *t*-mer position *i* in the window is considered by 𝒪_*k*_ for exactly one *k*-mer *X*_*p*_.

Let *S* be the starting positions of the minimal *t*-mers in *W*. Then mod-sampling samples the *k*-mer *X*_*p*_ at position *p* = min{*i* mod *w* | *i* ∈ *S*}. We want to show that *X*_*p*_ is the leftmost minimal *k*-mer according to 𝒪_*k*_. If *p* = *i* mod *w* and 𝒪_*t*_(*W* [*i*..*i* + *t*)) = *o*, then

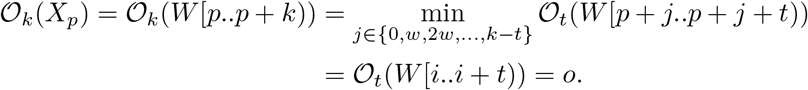

Since *o* is the minimal *t*-mer in the window, any other *k*-mer *X*_*q*_ must have order 𝒪_*k*_(*X*_*q*_) ≥ *o*. By choice of *p* as the minimal position for which 𝒪_*k*_(*X*_*p*_) = *o* holds, *X*_*p*_ is indeed the leftmost minimal *k*-mer according to 𝒪_*k*_.

Our first example, the *lr-minimizer*, is inspired by syncmers [4] and miniception [27], and corresponds to the choice *t* = *k* − *w*. By Lemma 13, mod-sampling yields a minimizer schemes for this choice of *t* because *t* ≡ *k* (mod *w*). As already mentioned, we must ensure that *t* is sufficiently large. Let therefore *r* ≥ 1 be a lower bound on *t*.

#### Definition 14

(`lr-minimizer`). *When k* ≥ *w* + *r, choosing t* = *k* − *w gives the lr-minimizer*. The name “lr” (short for “left-right”) comes from the fact that when the minimal *t*-mer is in position 0 ≤ *x < w*, the *t*-mer is a prefix of the sampled *k*-mer (so, they are left aligned), and when *w* ≤ *x <* 2*w*, the *t*-mer is a suffix of the sampled *k*-mer (so, they are right aligned). The lr-minimizer is related to both the miniception and the closed syncmer: miniception subsamples closed syncmers by a random order 𝒪_*k*_, whereas the lr-minimizer subsamples them based on the order 𝒪_*t*_ of the minimal contained *t*-mer, obtaining a lower density.

#### Corollary 15.

*The density of the lr-minimizer* 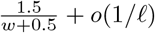.

**Proof**. Immediate by using *t* = *k* − *w* in Theorem 10 and using 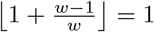.

Our second example gives the main result of this work, the mod-minimizer.

#### Definition 16

(mod-minimizer). *Let r be a (small) integer lower bound on t. For any k* ≥ *r, choosing t* = *r* + ((*k* − *r*) mod *w*) *gives the mod-minimizer*.

Compared to the lr-minimizer, the mod-minimizer uses a smaller *t* and the minimal *t*-mer of a window is not necessarily a prefix or a suffix of the sampled *k*-mer. Here, the name is inspired from the fact that we choose *t* using the mod function.

#### Corollary 17.

*The density of the mod-minimizer is* 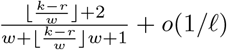 *which tends to* 1*/w for large k and r* = ⌈(3 + *ϵ*) log_*σ*_(*ℓ*)⌉.

**Proof**. Immediate from Lemma 9, Theorem 10, and Corollary 11.

In practice, choosing *r* slightly above log_*σ*_(*ℓ*) is sufficient to mitigate duplicate *t*-mers and achieve good density. For example, in our experiments we use *r* = 4.

We remark that Marçais et al. [11] were the first to describe a method that achieves optimal density when *k* → ∞, i.e., the “rotational” minimizer reviewed in Section 3. However, their approach is particularly involved as well as the proof of optimality. On the contrary, the mod-sampling algorithm is very intuitive (see also Figure 1) and the proof of optimality for the mod-minimizer is almost straightforward. Furthermore, as we are going to see in Section 5, the mod-minimizer performs better than the rotational minimizer for practical values of *k* and *w*, i.e. it converges faster to 1*/w* compared to the rotational minimizer.

Lastly, it is easy to see that the random minimizer from Definition 4 is obtained by setting *t* = *k* in mod-sampling. As expected, using *t* = *k* in Formula 1 gives density 2*/*(*w* + 1).

## 5 Experiments

In this section we compare our new minimizer schemes against the approaches reviewed in Section 3 to highlight their benefits and discuss potential limitations. For all experiments we use the C++ implementation of the algorithms, compiled with `gcc` 11.1.0 under Ubuntu We use a 128-bit (pseudo) random hash function [22] to implement a random order.

### Density

We measure the density of the schemes on a synthetic sequence of 10 million i.i.d. random characters. We use *σ* = 4 (e.g., the DNA alphabet) in all experiments to highlight the direct link to concrete applications. In practice, fixing *r* = 4 is sufficiently large, and we set *t* = *r* + ((*k* − *r*) mod *w*) for for miniception, lr-minimizer, and mod-minimizer. Figure 5 shows the density of the schemes by varying *k*, for two example values of *w*. For sufficiently large *k*, we consistently have the following order, from best to worst density: (1) mod-minimizer, (2) rot-minimizer (our own alternative version given in Algorithm 7 in the Appendix), (3) lr-minimizer^5^, (4) miniception, (5) random minimizer. These plots confirm the theoretical result of Corollary 11: as *k* increases, the mod-minimizer approaches the optimal density 1*/w*. In particular, the mod-minimizer approaches 1*/w* faster than the rot-minimizer by Marçais et al. [11]. The Mykkeltveit MDS based methods of Pellow et al. [17] only perform well for small *k*.

**Figure 5.**
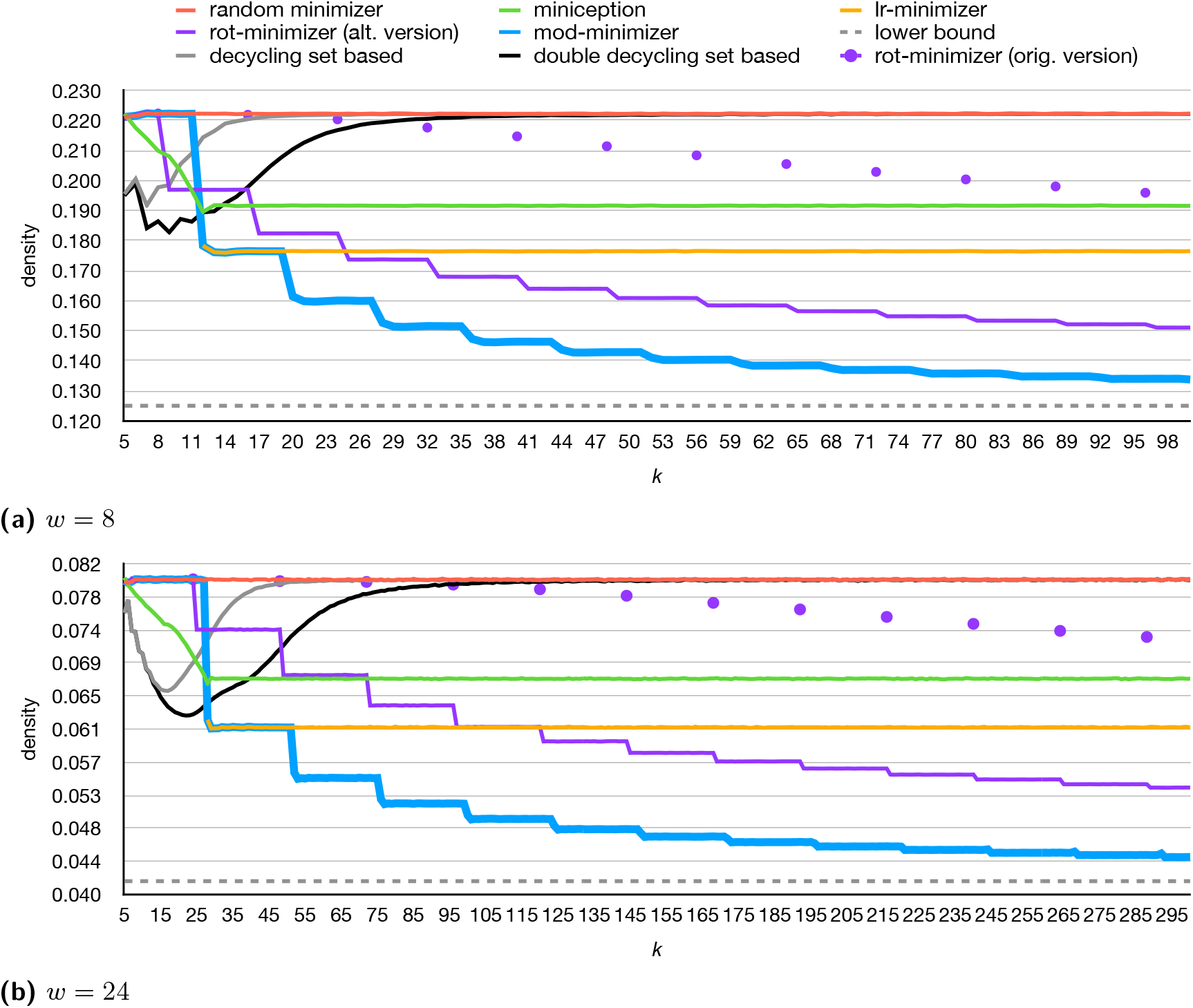
Density by varying *k* measured on a random string of 10 million characters for *σ* = 4. Miniception, lr-minimizer, and mod-minimizer all use *r* = 4 and *t* = *r* + ((*k* − *r*) mod *w*). The mod-minimizer has the same performance as the random minimizer when *k* ≤ *w* as *t* = *k* for such cases, whereas lr-minimizer requires *k* ≥ *w* + *r*. The original rot-minimizer is only defined when *w* divides *k*. For *w* = 24 and *k > σ*^*r*^ = 4^4^ = 256, the density of mod-minimizer has small spikes caused by duplicate *t*-mers.

### Sampling speed

As already noted in Section 3, all considered methods can (and, hence, *should* be used) in a *streaming* way, by taking advantage of overlapping computations across consecutive windows. Our implementation supports such sampling modality as well. All methods apart from the decycling methods perform essentially the same when streaming, taking around 30 to 40 nanoseconds per window on a Intel(R) Xeon(R) Platinum 8276L CPU clocked at 2.20 GHz, as long as *w, k* ≤ 100, where around half the time is spent on hashing *k*-mers (or *t*-mers). This is around 10× faster than sampling each window independently.

### Indexing *k*-mer sets with SSHash

As a concrete application of the mod-minimizer, we plug it into SSHash [18, 19], a recent *k*-mer dictionary based on minimizers and minimal perfect hashing [21]. In short, SSHash is a compressed data structure that maintains a set of *k*-mers (i.e., a De Bruijn graph) and supports exact membership queries. Minimizers are used as “seeds” to search the query *k*-mer into the data structure. Hence in this context, *k* is the total number of characters in a window (which we would otherwise write as *ℓ* = *w* + *k* − 1) of *w* consecutive *m*-mers, where *m* is the chosen minimizer length (otherwise *k* in this paper). The reference implementation of SSHash (at https://github.com/jermp/sshash) uses the random minimizer for simplicity and versatility. We replace it with the mod-minimizer and observe no slowdown in construction and query time, but better space usage. We report some example data.

We test default parameters *k* = 31 and *m* = 21. On human chromosome 13 random minimizers yield an SSHash index taking 7.53 bits/*k*-mer. Using mod-minimizers the space reduces to 6.41 bits/*k*-mer (−14.9%). On the whole human genome (GRCh38), space usage goes down from 8.70 to 7.40 bits/*k*-mer (−14.9%). On a small pangenome of 100 *S. Enterica* genomes [24], space usage goes down from 7.55 to 6.52 bits/*k*-mer (−13.6%). We also test one of the largest genome assemblies, the *Ambystoma Mexicanum* (the “axolotl”), containing over 18 billion distinct *k*-mers for *k* = 31. Again using *m* = 21, space goes down from 9.91 to 8.50 bits/*k*-mer (−14.2%).

Lastly, we point out the main limitation of the mod-minimizer. The mod-minimizer improves over random minimizers when *k > w*, or in SSHash’s notation, when *m >* (*k* + 1)/2. In practice, *m* is typically not larger than 21 because most minimizers already appear once for *m* = 21, while *k* can be up to 63. Thus, the mod-minimizer is only useful in specific cases.

## 6 Conclusions and future work

In this work, we introduced a simple framework, *mod-sampling*, that yields minimizer and local schemes, and has asymptotically optimal density for large *k*. Experimental results confirm our theoretical findings and demonstrate that a specific instantiation, the mod-minimizer, offers a good reduction in density compared to other methods. The mod-minimizer can be used as a drop-in replacement for random minimizers, for example, in the SSHash data structure. Our implementations are publicly available.

Future work could investigate the impact of different orders 𝒪_*t*_ for the *t*-mers. For example, combining the decycling set based order [17] with the mod-minimizer seems a promising idea. This could lead improved density when *k* ∼ *w*.

Contrary to the large-*k* case, for which there are asympotically optimal methods, the small-*k* case is not yet well understood. It is known that it is *not* possible to match the trivial lower bound of 1*/w* as Marçais et al. [11] give an improved lower bound. In Appendix A.2, we simplify their bound from 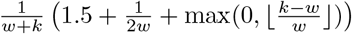 to 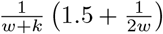, and show that it only improves over 1*/w* when *k <* (*w* + 1)*/*2. In Appendix A.3, we slightly improve the bound to 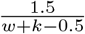 and extend it to hold for all *local* schemes. However, even in the simplest non-trivial case *k* = 1, *w* = 2, the best possible densities of schemes are only known for small alphabet sizes via exhaustive search, and are larger than the lower bound. Generally, it is not known whether Marçais’ lower bound is close to the true optimal density, although we conjecture it is not. Thus, the big open problem is to derive a tight lower bound on the optimal density of local/forward/minimizer schemes for small *k*.

## Algorithm 7

Pseudocode for the alternative implementation of the rotational minimizer algorithm that we also use in the experiments of Section 5.

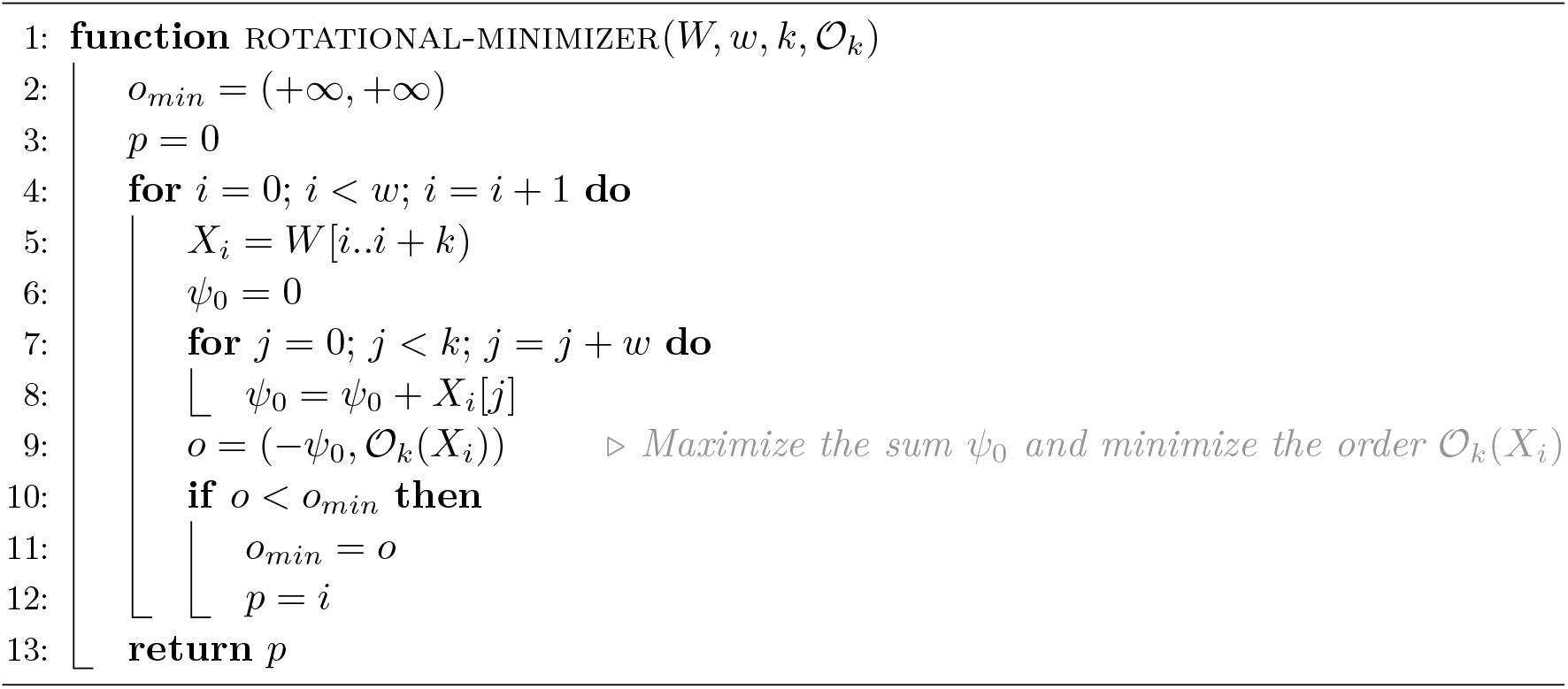

## A Appendix

### A.1 The probability of two duplicate *t*-mers in a window

We prove here the following lemma, adapted from Lemma 9 of [27].

#### Lemma 9.

*For any ϵ >* 0, *if t >* (3 + *ϵ*) log_*σ*_(*ℓ*), *the probability that a random window of ℓ* − *t* + 1 *t-mers contains two identical t-mers is o*(1*/ℓ*). *Given that ℓ* = *w* + *k* − 1, *o*(1*/ℓ*) → 0 *for large k*.

**Proof**. Following the proof of Lemma 9 of [27], we first bound the probability that two fixed *t*-mers are equal by *P* and then conclude that the probability that the window contains two identical *t*-mers is (*ℓ* − *t* + 1)^2^ · *P* by the union bound.

We distinguish two cases. (1) Assume the two fixed *t*-mers do not share any symbol. In this case, since the window is made of random symbols, the probability that they are the same is 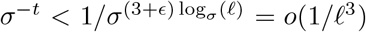. (2) Otherwise, they have an overlap. Call *x* the sub-string of the window made by the union of the two overlapping *t*-mers. When the two *t*-mers are offset by *d* positions, *x* has length *d* + *t* and, if the two *t*-mers are the same, *x* is a periodic string with period *d*. Hence, it is uniquely determined by its first *d* symbols and there are *σ*^*d*^ possible strings *x* for which the two *t*-mers are identical. The probability that a random *x* contains two identical *t*-mers is therefore *σ*^*d*^*/σ*^*d*+*t*^ = *σ*^−*t*^ = *o*(1*/ℓ*^3^). In conclusion, *P* = *o*(1*/ℓ*^3^) and the lemma follows via (*ℓ* − *t* + 1)^2^ · *P* ≤ *ℓ*^2^ · *P* = *ℓ*^2^ · *o*(1*/ℓ*^3^) = *o*(1*/ℓ*).

### A.2 Marçais et al.’s lower bound is only useful for small *k*

Marçais et al. [11] show the following lower bound for forward schemes:

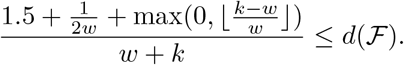

Assume that *k* ≥ 2*w*, so that the floor-term is non-zero. Write *k* = *c* · *w* with *c* ≥ 2. Then the lower bound is

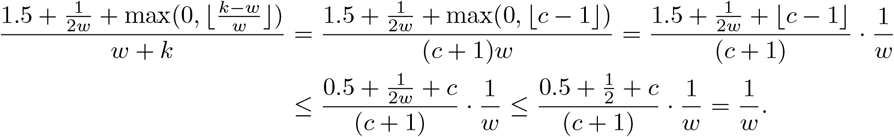

Thus, as soon as the floor term is non-zero, the lower bound is anyway less than the trivial lower bound of 1*/w*. Thus, we may as well use the simpler form

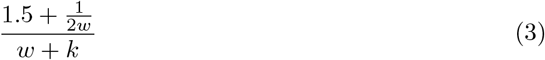

which closely mirrors the bound 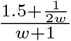 by Schleimer et al. [26]. Lastly, solving 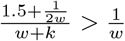 gives that Marçais et al.’s bound improves over the trivial lower bound for *k <* (*w* + 1)*/*2.

### A.3 Improving and extending Marçais et al.’s bound

Here we slightly improve the bound in Formula 3 and generalize it from forward schemes to local schemes.

#### Theorem 18

(New density lower bound). *The density of any local scheme satisfies*

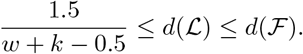

**Proof**. We assume the input is an infinitely long random string *S* over Σ. The proof makes use of the setting illustrated in Figure 6, which is as follows. We partition the windows of *S* in consecutive groups of 2*ℓ* + 1 windows. Let one such group of windows start at position *i*, and consider windows *W* and *W* ^*′*^ starting at positions *i* and *i*^*′*^:= *i* + *ℓ* respectively. Also let *W* ^*′′*^ be the window that is the exclusive end of the group, thus starting at position *i*^*′′*^ = *i* + 2*ℓ* + 1 = *i*^*′*^ + *ℓ* + 1. Note that there is no gap between the end of window *W* and the beginning of window *W* ^*′*^, whereas there is a gap of a single character between the end of *W* ^*′*^ and the beginning of *W* ^*′′*^ (shown as the gray shaded area in Figure 6). These three windows are disjoint and hence the random variables *X, X*^*′*^, and *X*^*′′*^ indicating *f*(*W*), *f*(*W* ^*′*^), and *f*(*W* ^*′′*^) respectively are independent and identically distributed. (But note that we do not assume they are uniformly distributed, as that depends on the choice of the sampling function *f* .) In Figure 6, we have *X* = 1 and *X*^*′*^ = *X*^*′′*^ = 2.

**Figure 6.**
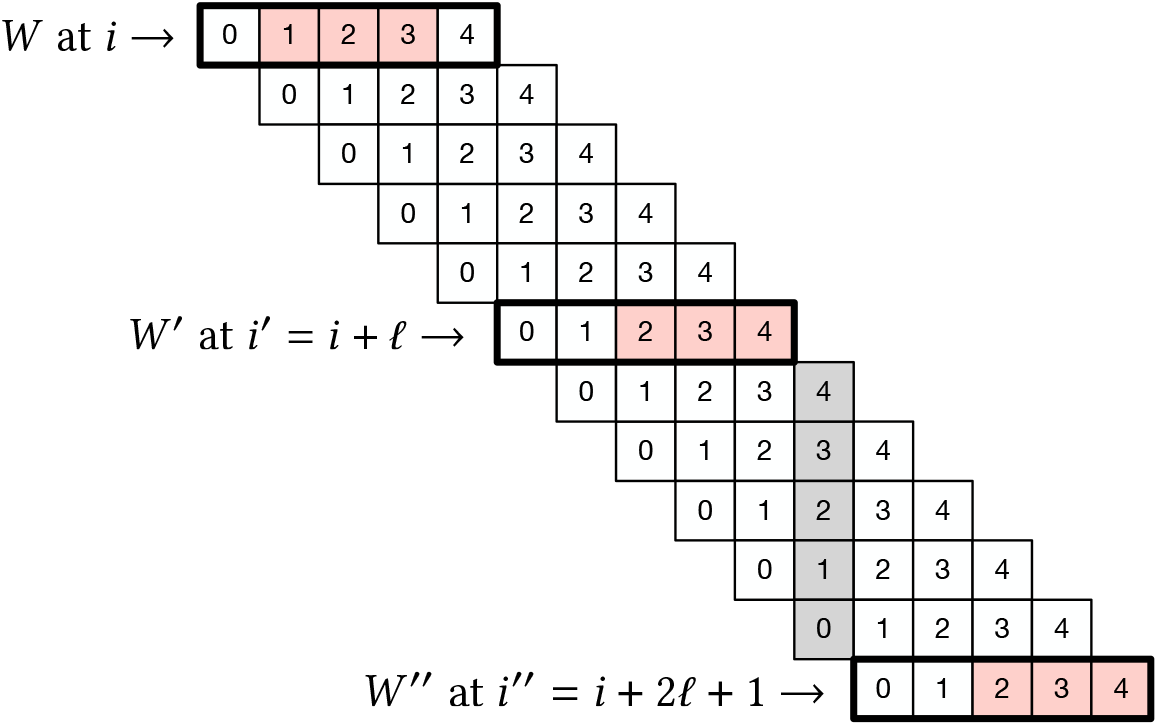
The lower bound setting from Theorem 18. In this example, we use *w* = *k* = 3, so *ℓ* = *w* + *k* − 1 = 5. Red boxes indicate the sampled *k*-mer in windows *W*, *W* ^*′*^, and *W* ^*′′*^ that are highlighted with a ticker stroke.

Since the three windows *W*, *W* ^*′*^, and *W* ^*′′*^ are disjoint, they sample *k*-mers at distinct positions (indicated by the red boxes in Figure 6). The proof consists in computing a lower bound on the expected number of sampled *k*-mers in the range [*i* + *X, i*^*′′*^ + *X*^*′′*^). Note that for non-forward schemes, it is possible that windows before *W* or after *W* ^*′′*^ sample a *k*-mer inside this range. For our lower bound, we will simply ignore such sampled *k*-mers.

When *X < X*^*′*^, the window starting at *i*+*X* + 1 ends at *i*+*X* +*ℓ* = *i*^*′*^ +*X < i*^*′*^ +*X*^*′*^, thus at least one additional *k*-mer must be sampled in the windows between *W* and *W* ^*′*^. Similarly, when *X*^*′*^ ≤ *X*^*′′*^, the window starting at *i*^*′*^ +*X*^*′*^ + 1 ends at *i*^*′*^ +*X*^*′*^ +*ℓ* = *i*^*′′*^ +*X*^*′*^ −1 *< i*^*′′*^ +*X*^*′′*^, so that at least another *k*-mer must be sampled in the windows between *W* ^*′*^ and *W* ^*′′*^.

Thus, the number of sampled *k*-mers in between positions *i* + *X* and *i*^*′′*^ + *X*^*′′*^ is at least 1 + ℙ[*X < X*^*′*^] + 1 + ℙ[*X*^*′*^ ≤ *X*^*′′*^]. Since *X, X*^*′*^, and *X*^*′′*^ are i.i.d., we have that ℙ[*X*^*′*^≤ *X*^*′′*^] = ℙ[*X*^*′*^ ≤ *X*] = 1 − ℙ[*X < X*^*′*^], and hence

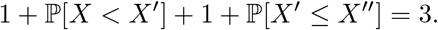

Since there are 2*ℓ* + 1 windows in each group, by linearity of expectation, we obtain density

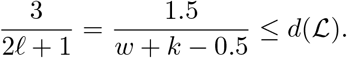

We point out that Schleimer et al.’s [26] lower bound on the density of “random” local schemes of 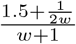 can be improved with the same technique to obtain a lower bound of 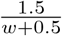, which coincidentally happens to exactly match the density of the lr-minimizer (Corollary 15). Note also the similarity of the latter bound with that of Theorem 18: Schleimer et al. assume independent hashes and obtain a denominator of *w* + 0.5, while Marçais et al. do not and hence *k* − 1 surrounding positions of context can be used, leading to *w* + *k* − 0.5.

To conclude, Figure 7 gives a visual comparison between the lower bounds considered here. Note that both bounds are only relevant for *k <* (*w* + 1)*/*2 as they are less than 1*/w* otherwise. Also, by solving 1.5*/*(*w* + *k* −0.5) *>* (1.5 + 1*/*(2*w*))*/*(*w* + *k*), we again determine that the improved bound is better than the simplified one exactly when *k <* (*w* + 1)*/*2.

**Figure 7.**
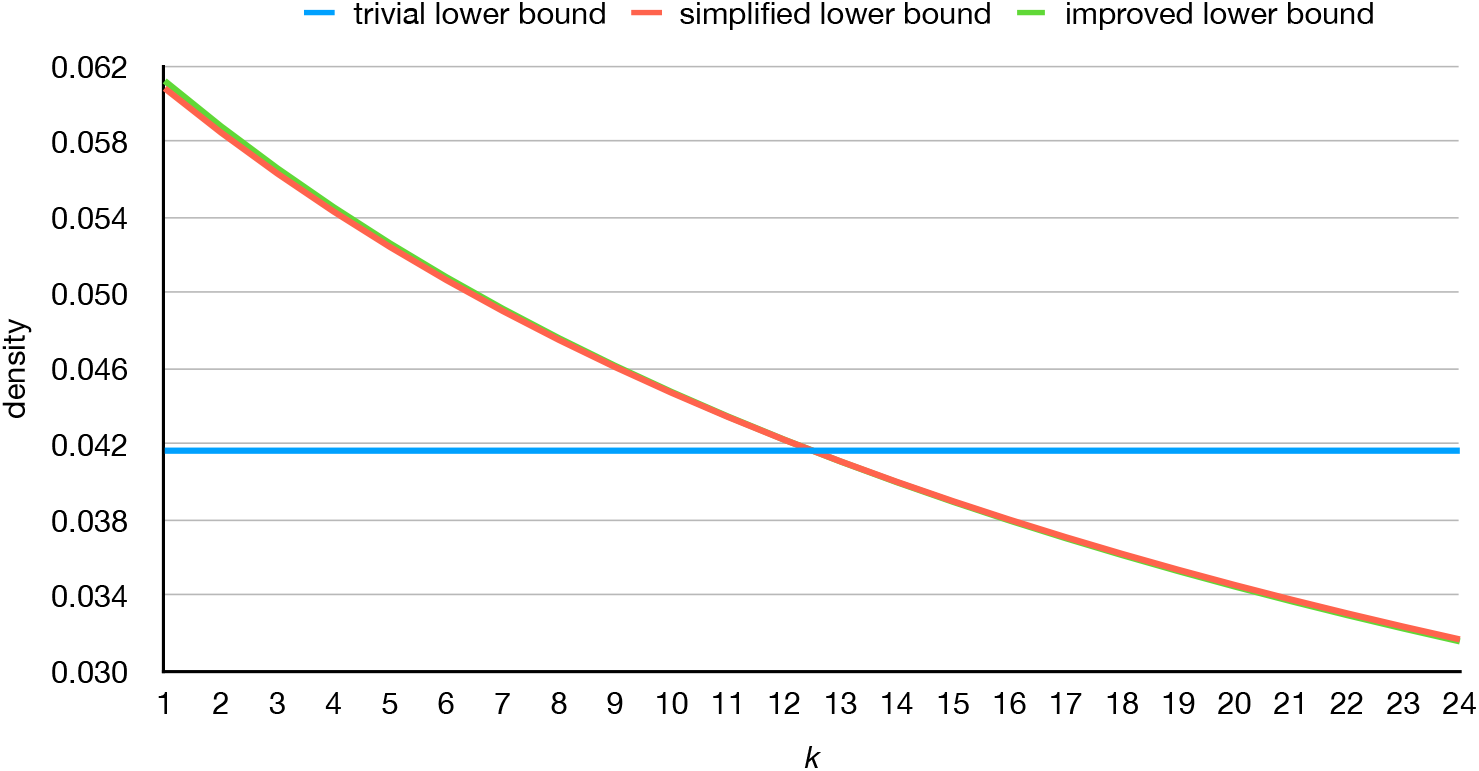
Comparison between the trivial 1*/w* lower bound, the simplified Marçais et al. [11] lower bound from Appendix A.2, and the new improved bound from Theorem 18, for *w* = 24.

### A.4 The shape of the density function in Formula 1

Here we prove the following lemma, as stated at page 11, and re-stated here for simplicity.

#### Lemma 12.

To ease notation, in the following we refer to the density of mod-sampling (Formula 1) as

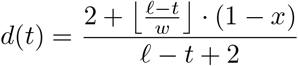

where *x* = 1*/*(*ℓ* − *t* + 1) if *t*≢ *k* (mod *w*), and *x* = 0 otherwise. To prove the lemma, we first make the following observations.

#### Observation 19.

*Let* 0 ≤ *q* ≤ ⌊*k/w*⌋ *be a given integer. For*

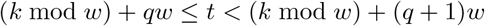

*the quantity* 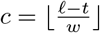 *is constant. Moreover*, 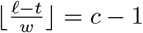 *for t* = (*k* mod *w*) + (*q* + 1)*w*.

**Proof**. Let *t*_*i*_:= (*k* mod *w*) + *qw* + *i*, for 0 ≤ *i < w*. Then

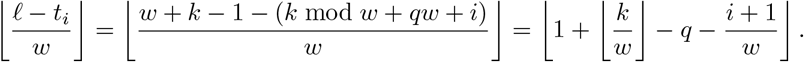

Since 0 ≤ *i < w*, we have 1*/w* ≤ (*i* + 1)*/w* ≤ 1, and hence the quantity remains constant. It is also easy to see that 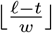 becomes *c* − 1 when *t* = (*k* mod *w*) + (*q* + 1)*w*.

#### Observation 20.

*Let* 0 ≤ *q* ≤ ⌊*k/w*⌋ *be a given integer. For*

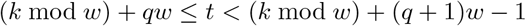

*d*(*t*) *is increasing, i*.*e*., *d*(*t*) *< d*(*t* + 1).

**Proof**. First, let us consider the case (*k* mod *w*) + *qw < t <* (*k* mod *w*) + (*q* + 1)*w* − 1. Then *t*≢ *k* (mod *w*), and the value of *x* in Equation (2) is equal to 1*/*(*ℓ* − *t* + 1). From Observation 19, 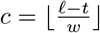 is constant. Hence, after quite some (omitted) steps, *d*(*t*) *< d*(*t*+1) simplifies to 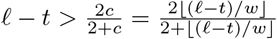 Now, let *y* = *ℓ* − *t* ≥ *w* −1 *>* 0 so that the previous inequality can be rewritten as 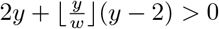. We know that 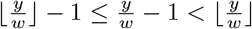, hence 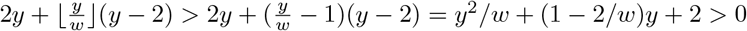 which is always true because *w* ≥ 2. Now consider the case *t* = (*k* mod *w*) + *qw*. In this case we have that the value of *x* in Equation (2) is 0 for *d*(*t*) and *d*(*t*) *< d*(*t* + 1) eventually simplifies to 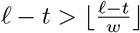, which is again always true because *w* ≥ 2.

#### Observation 21.

*For any* 0 ≤ *q* ≤ ⌊*k/w*⌋, *d*((*k* mod *w*) + *qw* − 1) ≥ *d*((*k* mod *w*) + *qw*).

**Proof**. In this case, set *t* = (*k* mod *w*) + *qw* − 1. We have *t* + 1 ≡ *k* (mod *w*) so that *x* = 0 for *d*(*t*+ 1), and obtain 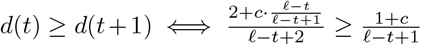 where 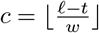 is a constant (Observation 19) and this eventually simplifies to 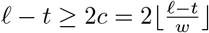, which is true because *w* ≥ 2.

#### Observation 22

*For any* 0 ≤ *q* ≤ ⌊*k/w*⌋, *d*((*k* mod *w*) + *qw*) *< d*((*k* mod *w*) + (*q* + 1)*w*).

**Proof**. Let *t* = (*k* mod *w*) + *qw*. Clearly, *t* ≡ *k* (mod *w*) and (*t* + *w*) ≡ *k* (mod *w*), hence *x* = 0 in Equation (2). So 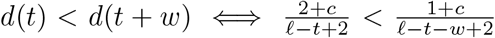 where 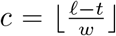 is a positive constant for Observation 19. This eventually simplifies to 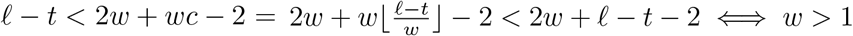.

From these observations, Lemma 12 follows: the density function from Theorem 10 has local minima at *t* ≡ *k* (mod *w*) achieves its minimum on *r* ≤ *t* ≤ *k* either at the domain boundary *t* = *r*, or the smallest multiple of *w* in the domain, *r* + ((*k* − *r*) mod *w*). See Figure 4 at page 11 for some concrete examples.

## Footnotes

1 In practice, the density ≈ 2*/*(*w* + 1) is achieved even when 𝒪_*k*_ is not *total*, but simply when 𝒪_*k*_ is a hash function with a sufficiently large output range that avoids collisions among the *w k*-mers in a window.

2 Guillaume Marçais approved the name (personal communication).

3 Pellow et al. [17] consistently write *counterclockwise*, but both their code and example figure, as well as Mykkeltveit’s original definition [14], use the first *clockwise* rotation with positive imaginary part. Indeed, by “counterclockwise”, they refer to the left cyclic rotation of a *k*-mer and not to the rotation of the complex number (personal communication).

4 Or, practically, take the value of *t* that minimizes Formula 1, regardless of whether *t* ≡ *k* (mod *w*) or not.

5 Miniception and lr-minimizer are context-sensitive versions of the closed syncmer [4], and thus we omit closed syncmers in Figure 5.

